# MELODI - Mining Enriched Literature Objects to Derive Intermediates

**DOI:** 10.1101/118513

**Authors:** Benjamin Elsworth, Karen Dawe, Emma E Vincent, Ryan Langdon, Brigid M Lynch, Richard M Martin, Caroline Relton, Julian PT Higgins, Tom R Gaunt

## Abstract

**Motivation:** The scientific literature contains a wealth of information from different fields on potential disease mechanisms. However, prioritising mechanisms for further analytical evaluation presents enormous challenges in terms of the quantity and diversity of published research. The application of data mining approaches to the literature offers the potential to identify and prioritise mechanisms for more focused and detailed analysis.

**Results:** Here we present MELODI, a literature mining platform that can identify mechanistic pathways between any two biomedical concepts. Two case studies demonstrate the potential uses of MELODI and how it can generate hypotheses for further investigation. Firstly, an analysis of ERG and prostate cancer derives the intermediate transcription factor SP1, recently confirmed to be physically interacting with ERG. Secondly, examining the relationship between a new potential risk factor for pancreatic cancer identifies possible mechanistic insights which can be studied *in vitro.*

**Availability:** MELODI has been implemented as a Python/Django web application, and is freely available to use at www.melodi.biocompute.org.uk

## 1 Introduction

An understanding of the mechanisms that relate risk factors to health outcomes is important in the discovery of potential drug targets and disease biomarkers. Identifying the mechanistic pathway from a given exposure to a given disease allows us to consider which steps along the pathway are potentially modifiable. This offers the potential to identify new biomarkers and potential treatments to reduce subsequent risk of the disease.

Before embarking on novel research, either *in vitro* or *in silico*, it is important to examine what evidence is already available, so that the most promising opportunities are pursued and resources are not wasted. However, most scientific subjects produce too many publications to read in detail, and consequently researchers often rely on inefficient and potentially biased methods to prioritise which mechanisms to investigate. Existing approaches to collating relevant published evidence include manual filtering/selection of the literature, ranking by publication metrics such as impact factor / number of citations, examining media/social media reports and word of mouth. It is clear that a more systematic, automated data mining approach offers enormous potential to assist in identification of existing evidence and therefore prioritisation of mechanisms to investigate. To this end, tools which help us search and refine a set of literature are becoming increasingly important.

To derive potential mechanisms on the pathway between two concepts we can search the literature for overlapping mechanisms between these two concepts. One option to do this is to simply search the two concepts simultaneously with a boolean ‘AND’ operator, using PubMed (http://www.ncbi.nlm.nih.gov/pubmed) or a standard search engine. This approach, however, will only find cases where both concepts are described together in the same place. This will miss cases where the overlapping mechanisms are described in independent places, i.e. in one place in relation to the risk factor and in another in relation to the outcome. To address this we need to assess each concept separately and look for overlapping elements.

Many text mining tools are available that can be used to extract potential mechanistic terms from the free text of articles, usually titles and abstracts. A search of OMICtools (March 2017) produced 104 tools in the ‘Information Extraction’ category alone (http://omictools.com/information-extraction-category). These vary widely in their approach to key aspects such as synonyms, performance and precision. An alternative to extracting information from raw literature text is to use pre-calculated literature annotation objects. These can either be generated by humans, e.g. Medical Subject Headings (MeSH) (http://www.nlm.nih.gov/mesh/) or computationally derived, e.g. Semantic MEDLINE Database (SemMedDB) (Kilicoglu *et al.*, 2012).

The MeSH thesaurus is a well established set of terms used to index articles from over 5,000 of the world’s leading biomedical journals. These terms are freely provided by the National Library of Medicine and currently consist of over 27,000 descriptors, which are used by the MeSH staff to assign the most appropriate terms to each MEDLINE/PubMed article. Data are arranged in a complex hierarchy and are available to download in many different formats including summary data, hierarchy, and frequency.

SemMedDB is arepository of semantic predications extracted by SemRep (Rindflesch and Fiszman, 2003) which utilises the Unified Medical Language System (UMLS). For every title and abstract a Subject-PREDICATE-object triple is generated where the subject and object are terms from the UMLS Metathesaurus and the predicate is a relation from the UMLS Semantic Network. For example, the sentence "We used hemofiltration to treat a patient with digoxin overdose that was complicated by refractory hyperkalemia" produces the following four triples (http://semrep.nlm.nih.gov/):

1. Hemofiltration-TREATS-Patients
2. Digoxin overdose-PROCESS_OF-Patients
3. hyperkalemia-COMPLICATES-Digoxin overdose
4. Hemofiltration-TREATS(INFER)-Digoxin overdose

Identifying patterns and overlapping elements across two sets of scientific literature could involve either finding single terms common to both articles, e.g. MeSH or SemMedDB terms, or more complex associations, e.g. finding pairs of SemMedDB triples where the object of a triple in one article set is the subject of a triple in the other (Figure 2). This has been attempted by a number of others. One well-established method is Arrowsmith (Smalheiser *et al*., 2009), which is based on single overlapping words from article titles only,limited to a maximum of 25,000 articles per article set and has limited options to download, visualise, filter and explore the resulting data (Supplementary table 2). Another application is TeMMPo (https://www.temmpo.org.uk/) which was developed for identifying intermediate biological mechanisms underpinning the effects of lifestyle factors on cancer outcomes, using a predefined set of possible intermediates. This approach, however, is hypothesis driven (because intermediates have to be identified in advance) which limits the capacity to identify novel intermediates. There are also examples of searching SemMedDB data for overlapping elements (Hristovski *et al.*, 2015a; Cameron *et al.*, 2015) and even proof of principle examples of using graph databases (Hristovski *et al.*, 2015b). However, there is currently no modern software tool to perform this task on custom data sets in an efficient and intuitive way using the enrichment of annotation objects to refine the list of terms.

We present MELODI (Mining Enriched Literature Objects to Derive Intermediates), a web application which provides a user-friendly environment in which to identify overlapping intermediates from two sets of articles representing two distinct biomedical concepts. MELODI includes an enrichment step, whereby the frequencies of terms within a set of articles are compared with the background frequencies in the whole database. This step is particularly important for identification of overlapping elements due to the abundance of common terms and the error rate associated with extracting terms from free text. Enrichment prioritises cases where an object has been derived from text on multiple occasions, and often these are from independent studies, reducing the effects of incorrect assignment. MELODI also has an upper limit on the number of articles per article set of one million and via authentication provides user space which retains user specific search results and article sets.

## 2 Implementation

### 2.1 Application construction

We constructed MELODI using the Django web framework(http://djangoproject.com) with the following additional plugins and features. Authentication is handled via the Django Social Authenticationplugin (http://django-social-auth.readthedocs.org/en/latest/), providing a method to make all data, jobs and results both user specific and retrievable. As some of the database queries were proving too intensive for a responsive user experience, we integrated a task management system. This utilises thedistributed task queueCelery (http://www.celeryproject.org/) and the in-memory data structure storeRedis (http://redis.io/). Allowing jobs to be handled by a sophisticated task management system removes the need to wait on screen for analyses to complete and provides the opportunity for large complex queries. Supplementary figure 1 summarises the flow of the application.

### 2.2 The graph database

Identifying connections between two sets of articles can involve searching many millions of objects and relationships. Data storage and analysis of this type is suited to graph databases, and with recent advances in this field we decided from the outset to use a graph database (Neo4j,http://neo4j.com/). Neo4j is a database constructed of nodes, relationships and properties which structures data on the basis of relationships rather than in a conventional tabular structure. We chose this approach in preference to a relational database due to the data being relationship rich, the predicted search strategies (i.e. identifying novel relationships between datasets) and the intuitive nature of using a graph to contain and search data of this type. We implemented an additional MySQLdatabase (http://www.mysql.com/) to provide job progress reports, record user specific job data and improve data processing at the front end.

### 2.3 Data

We preloaded the graph with publicly available data from MeSH and SemMedDB then augmented and modified with frequencies per year for each annotated term and user provided relationship data (Table 1). As of March 2017 the graph consists of over 44 million nodes and 200 million relationships, the main components of which are listed in table 2. We will update the graph as new releases of MeSH and SemMedDB data are released. Figure 1 displays a simplified version of a very small article set containing three articles.

**Fig. 1.**
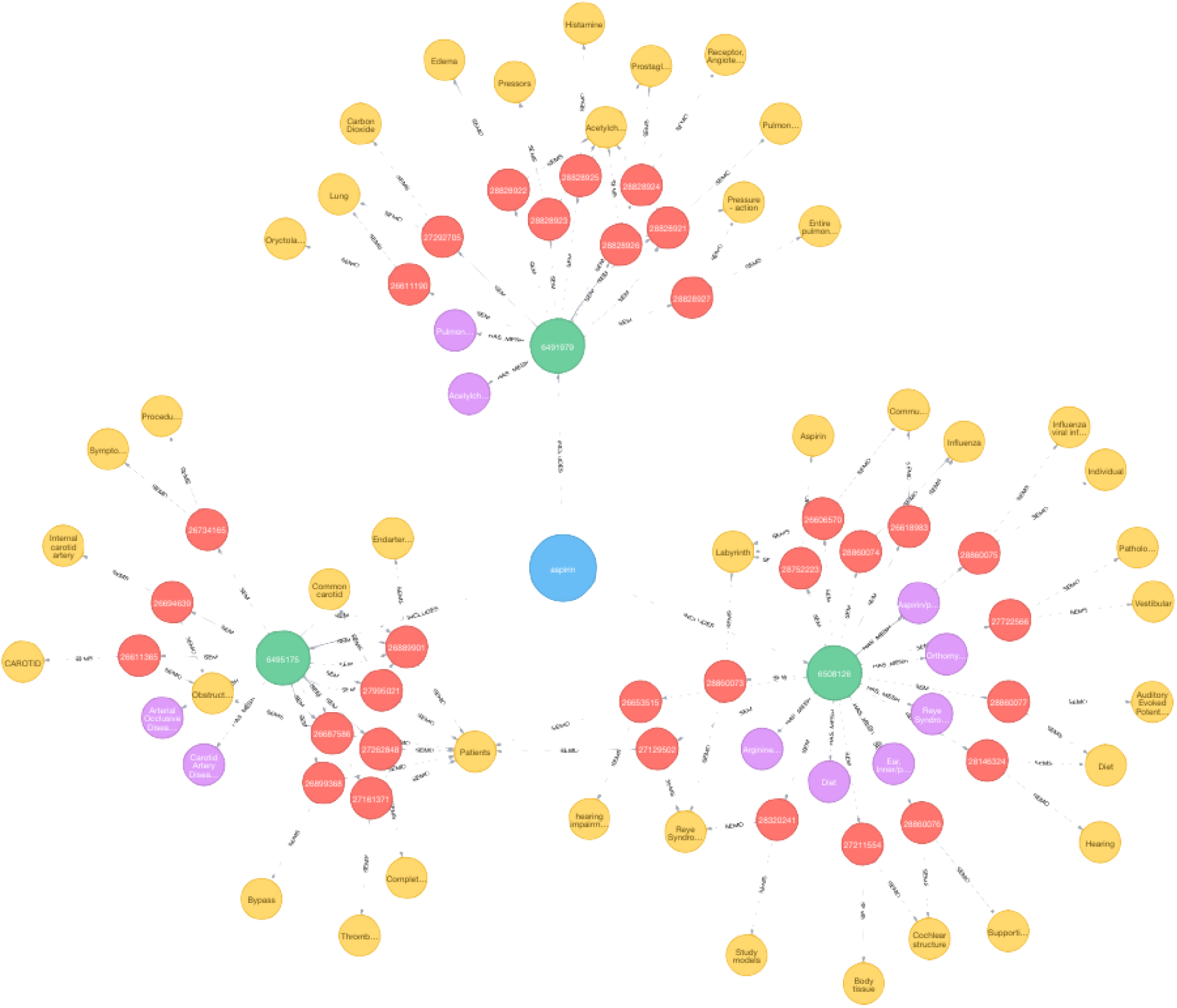
A three article ‘Article Set’. Each article set contains relationships to a set number of articles (green nodes) each of which has pre-defined relationships with MeSH/SemMedDB objects. Purple nodes represent MeSH terms, red and yellow the SemMedDB triples and concepts respectively.

**Table 1.** Sources of data

| Entry | Name | Version | Description |
| --- | --- | --- | --- |
| P | MeSH counts* | 2015 | Frequency counts for main MeSH terms |
| P | MeSH structure** | 2016 | MeSH hierarchy structure |
| P | PubMed*** | 26 | Basic article information up to 30/04/16 |
| P | PubMed-MeSH*** | 2016 | MeSH terms for each PubMed article |
| P | SemMedDB**** | 26 | SemMed summary data for each article |
| U | Article Set | N/A | Collection of PubMed articles |
| U | SearchSet-PubMed | N/A | Article set to PubMed relationships |
| C | New PubMed | Daily | PubMed information not already in database |
| C | PubMed-Mesh | Daily | PubMed to MeSH relationships |

P = Preloaded, U = User Uploaded, C = Computationally Uploaded
\* https://mbr.nlm.nih.gov/Download/
\*\* https://www.nlm.nih.gov/mesh/download_mesh.html
\*\*\* https://mbr.nlm.nih.gov/MRCOC.shtml
\*\*\*\* http://skr3.nlm.nih.gov/SemMedDB/index.html

**Table 2.** Graph details (Mar 2017)

| Data | Type | Number |
| --- | --- | --- |
| PubMed | Node | 25,698,930 |
| Mesh | Node | 464,122 |
| MeshTree | Node | 56,326 |
| SemMedDB Triple | Node | 17,713,740 |
| SemMedDB Concept | Node | 284,806 |
| PubMed - Mesh | Relationship | 80,395,022 |
| PubMed - SemMedDB Triple | Relationship | 84,621,296 |
| SemMedDB Triple - SemMedDB Concept | Relationship | 35,498,324 |

#### 2.3.1 Preloaded data

Table 1 lists the sources of data that were preloaded into the graph, notably summary data from SemMedDB and MeSH. We first transformed SemMedDB data sets from SQL and then all data were converted to a standard delimited format. We then inserted the data into the graph using the ‘neo4j-import’ command, a very efficient method for inserting large amounts of data into a Neo4j graph. Data needed to be pre-processed to include the same separator, correct header information and care taken so that all other insertion requirements were met. The application code, scripts that we used to transform theraw data into appropriate files and the neo4j import command are available at http://github.com/MRCIEU/melodi/. Once data were inserted, the graph was indexed in a similar way to standard relational databases.

#### 2.3.2 Uploaded data

Our main aim of preloading a large amount of data into the graph was to minimise the time spent processing user-supplied information. Therefore, the only modifications to the graph required when generating a user set of articles are the specification of the article set as a node and specification of the relationships between that node and the publications to which it relates, e.g. the lines connecting the blue ‘Article Set’ node and the green ‘Publication’ nodes in Figure 1. We initially attempted to create an empty graph that was organically populated with user information; however, the overheads associated with dynamically creating a graph using large amounts of data and using the less efficient insertion methods were too great.

### 2.4 Using the application

#### 2.4.1 Creating an article set

MELODI is based on identifying mechanisms linking two article sets. An article set is simply a set of articles that represent a defined concept, e.g. ‘body mass index’. There are two ways to create a new article set, each of which is defined by a set of PubMed IDs. The first is to perform a PubMed search within the application. This retrieves the PubMed IDs using the Entrez programming utilities (Sayers, 2009) and populates the graph with the new relationships as described above. The second option is to upload a set of PubMed IDs. The benefit of this option is that a set of IDs can be hand curated, improving precision and focus, removing the reliance on the methodology of a PubMed search and allowing for greater flexibility in how the set is created. When carefully constructed, producing article sets in this way will generate the most informative results with higher specificity (although this ‘manual curation’ has the potential to introduce additional bias).

#### 2.4.2 Comparing article sets

A simple hypothesis free approach to comparing the two article sets would be to identify terms which overlap across the two sets and quantify their occurrence using the number of articles mentioning them. This would identify the most common elements present in the article sets, but many of these will be uninformative (eg commonly occurring terms). MELODI reduces the problem with a two step strategy (Figure 2), using enrichment to identify terms that occur in the article sets more often than chance.

**Fig. 2.**
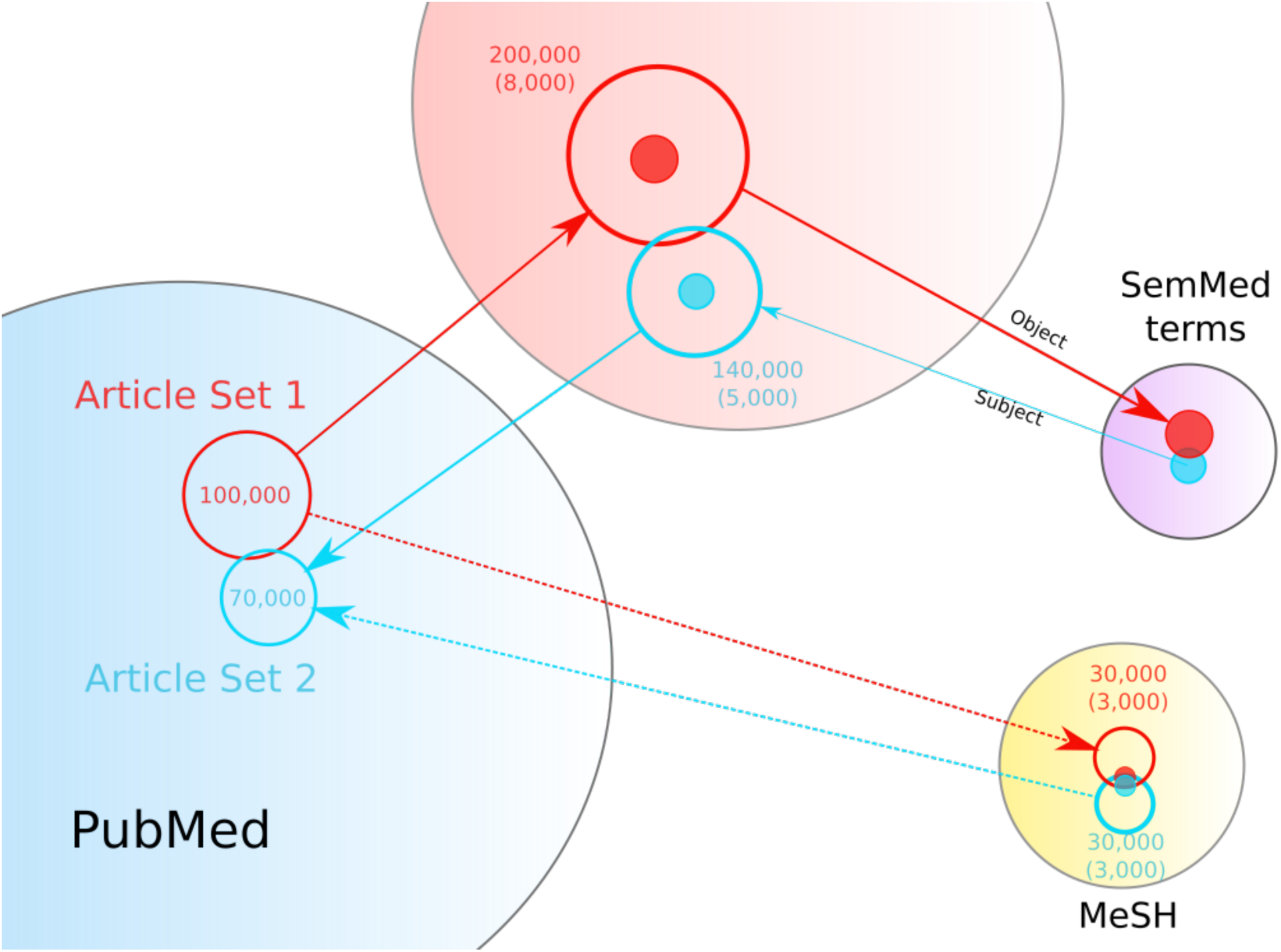
An overview of the enrichment analysis.An example analysis of two article sets, one with 70,000 articles (blue) and one with 100,000 articles (red). For each article set the SemMedDB and MeSH data are retrieved from the graph (open circles) with roughly twice as many objects as articles. An enrichments step identifies those objects that are enriched in that particular data set compared to the background (solid circles and numbers in parentheses). Those objects that are found to be overlapping between the article sets are returned for further analysis.

In the first step, for a given article set the enriched elements are identified. For MeSH terms, this is based on the number of times a MeSH term has been annotated as a main MeSH term in the article set, compared with the frequency of the main MeSH terms across all articles in MEDLINE (calculated from the entire set of MeSH data). For SemMedDB, two alternative analysis methods are available. The first is very similar to the MeSH approach, using the single SemMedDB terms (extracted from the triples) and then identifying enriched terms using their pre-calculated global frequencies. The second has a similar enrichment step but using the entire triplet and again frequencies calculated from the global dataset as the reference. An extra filtering stepis also employed for SemMedDB triples which restricts the objects of article set A and the subjects of article set B to terms which are present fewer than 150,000 times in the SemMedDB data set. This vastly reduces the number of overlapping terms, removing 88 high level terms such as ‘Patients’ and ‘Cells’ which would otherwise introduce unnecessary noise (Supplementary Table 1). In all three cases, the frequency of the term is compared between the article set and the entire literature set using a two-tailed Fisher’s exact test (FET). P-values are corrected for multiple testing using the Benjamini/Hochberg (non-negative) correction with a cutoff of p<1e-5. The results (p-value, corrected p-value and odds ratio) are stored on file for later use to avoid the need for repeating this step which can be computationally demanding. The enrichment ensures that when large numbers of articles are involved, multiple instances of a term are required to define a term as enriched, i.e. a single instance of a term is unlikely to be sufficient to be marked as enriched unless the article set in question is relatively small.

#### 2.4.3 Exploring the results

An analysis of two article sets can be performed using the MeSH terms and/or the SemMedDB data. The results of these independent approaches are provided separately for further investigation. As the analysis is hypothesis free we believe it is important to provide a first-pass filtered set of results on the initial view but also include access to the entire set of results, allowing the user to explore the data in its entirety. We have provided the following filtering parameters: FET corrected p-value, FET odds ratio and the top number of results (all methods), as well as minimum position in the MeSH hierarchy (MeSH method) and frequency of the predicate term (SemMedDB triple method). The inclusion of all predicates in the final data is deliberate, as cases where there are low numbers of enriched overlapping SemMedDB triple objects can still hold valuable information but may use more common predicates such as ASSOCIATED_WITH and AFFECTS. This is unlike otherapproaches (Cameron *et al.*, 2015) in which a universal predicate filter was used. As article set comparisons produce differing numbers of overlapping elements a basic set of dynamic filtering rules are employed when displaying the resultsfor the first time, adjusting the filters based on the number of results, e.g. low numbers of overlapping enriched elements are treated with a more relaxed set of parameters.

Visualisation of results is provided using interactive Sankey diagrams and for SemMedDB triples an additional force directed network diagram (Figure 3). Both are based on the same data displayed in the table on the page, with strength of supporting evidence represented by the thickness of connections. The network based view is often more intuitive when trying to identify paths between two article sets, as the same SemMedDB object may be present multiple times. This view also highlights the possibility of multiple steps between article sets.

**Fig. 3.**
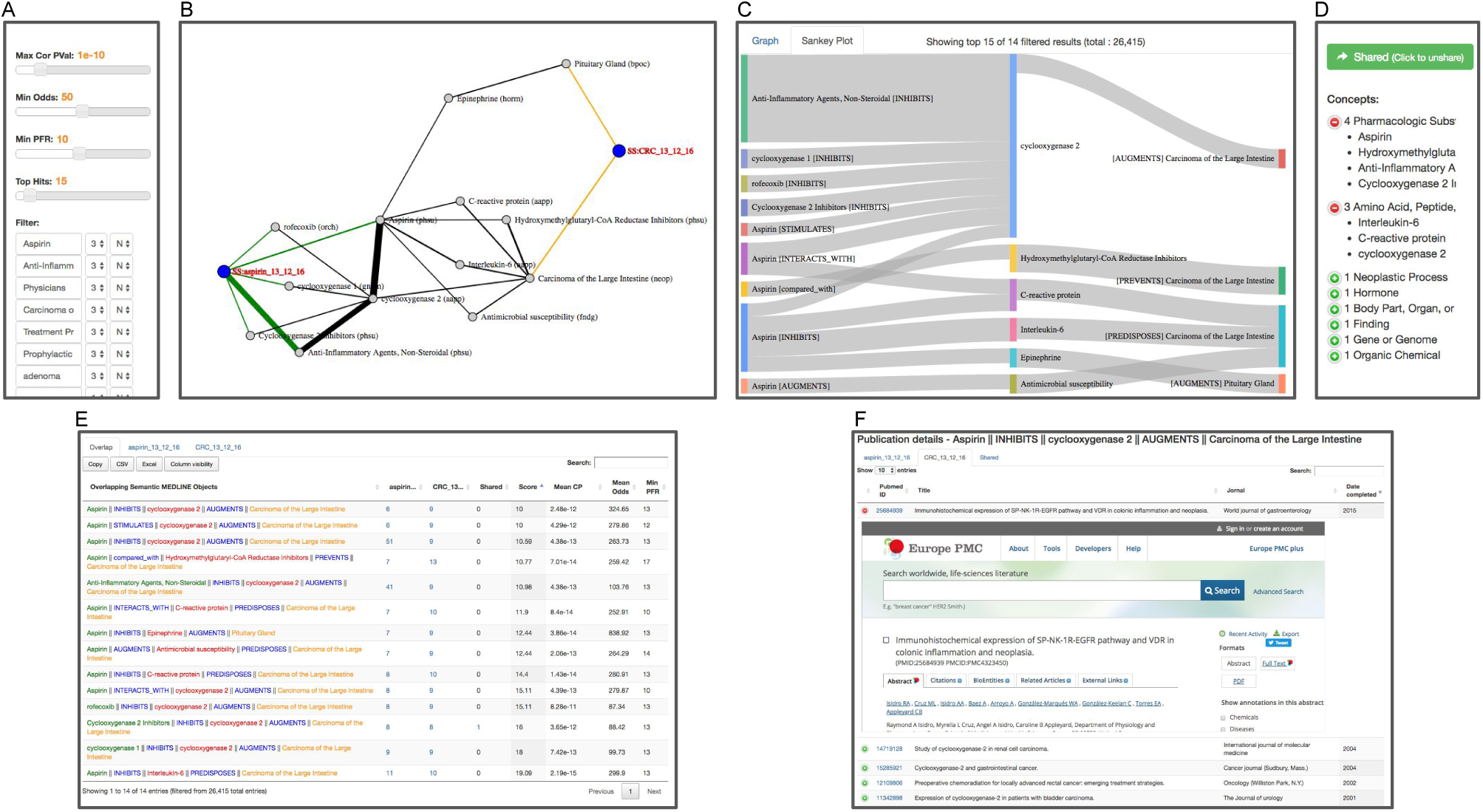
Exploring the results.An example of the visualisations and hlters within the results. Panel A contains the main filtering options for the Semantic Medline data, including P-value, odds ratio and Predicate Frequency Rank (PFR). This final metric allows the user to filter results based on the global frequencies of the SemMedDB predicates on the assumption that less frequent predicates are more informative. Panel B gives an example of the force directed graph visualisation of the results. Each article set (blue), subject and object (grey) are displayed as nodes with the relationships between them collapsed into single arcs. The thickness of these arcs represents the number of publications generating the relationship. Panel C represents the same data in a Sankey diagram, this time the individual predicates are listed in square brackets and the thickness of the bands represents the number of publications. Each SemMed subject and object as a type associated with it. Those present in the figures are listed in panel D. This gives an immediate overview of the types of terms present in the results on display. Panel E shows the table displayed below panels A-D on the results page. This table is the data source behind the visualisations, and therefore controls what is displayed. Clicking on any of the coloured terms adds that term to a hlter box in panel A which can then be set to exclude (N) or include (Y) that term in the results. The blue numbers link to the publications that derive the results. Panel F shows the detailed publication data from one of these links allowing the user to correlate the enriched concepts to the actual articles they were derived from.

The automatic filtering performed before visualisation attempts to highlight the most informative enriched overlapping elements. However, as free text is inherently noisy and computational predictions are not perfect, an additional text filtering option is provided. This consists of a number of text boxes that can be used either to ‘filter out’ or to ‘restrict to’ key words within the results, and in the case of SemMedDB analysis this can be done on any of the five elements (subject A - predicate 1 - object Alsubject B - predicate 2 - subject B) simply by clicking on the corresponding term in the table (Figure 3). The filter option is particularly useful where there are many overlapping terms between two article sets, and the restrict option is useful in cases where a more focused search is required, for example with defined exposure and outcome terms (Supplementary Figure 3). Examples of this are discussed later.

The order in which the results are delivered is critical, as this determines the top set that are displayed for closer inspection. This order can be based on a number of factors, e.g. the mean corrected p-value across the two article sets, the minimum position in the MeSH hierarchy or thepredicate frequency rank. In addition to these, a custom designed score is used which aims to identify overlapping elements with a high number of supporting articles in both article sets (Equation 1) where *uniq_a* and *uniq_b* are the number of unique articles from each article set that contain the overlapping term. This is based on the assumption that these elements may be the most reliable, due to many and equal articles producing the objects. This assumption does risk ignoring cases where a small number of valid articles support one side of the relationship and should therefore be used with caution.

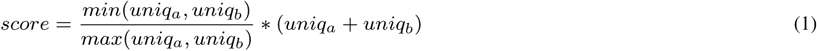

### 2.5 Applying MELODI to scientific research

The hypothesis generation methods provided by MELODI can be applied to a wide range of scientific disciplines. As we have implemented MELODI using PubMed and SemMedDB the current applications have a biomedical focus. In all cases the overarching aim would be to derive intermediates between two clearly defined scientific concepts. These two scientific concepts can arise from a range of different techniques and approaches, and the intermediate mechanisms identified can be further investigated using a similar range of techniques and approaches (Figure 4). For simplicity we have grouped some of these approaches into three categories.

**Fig. 4.**
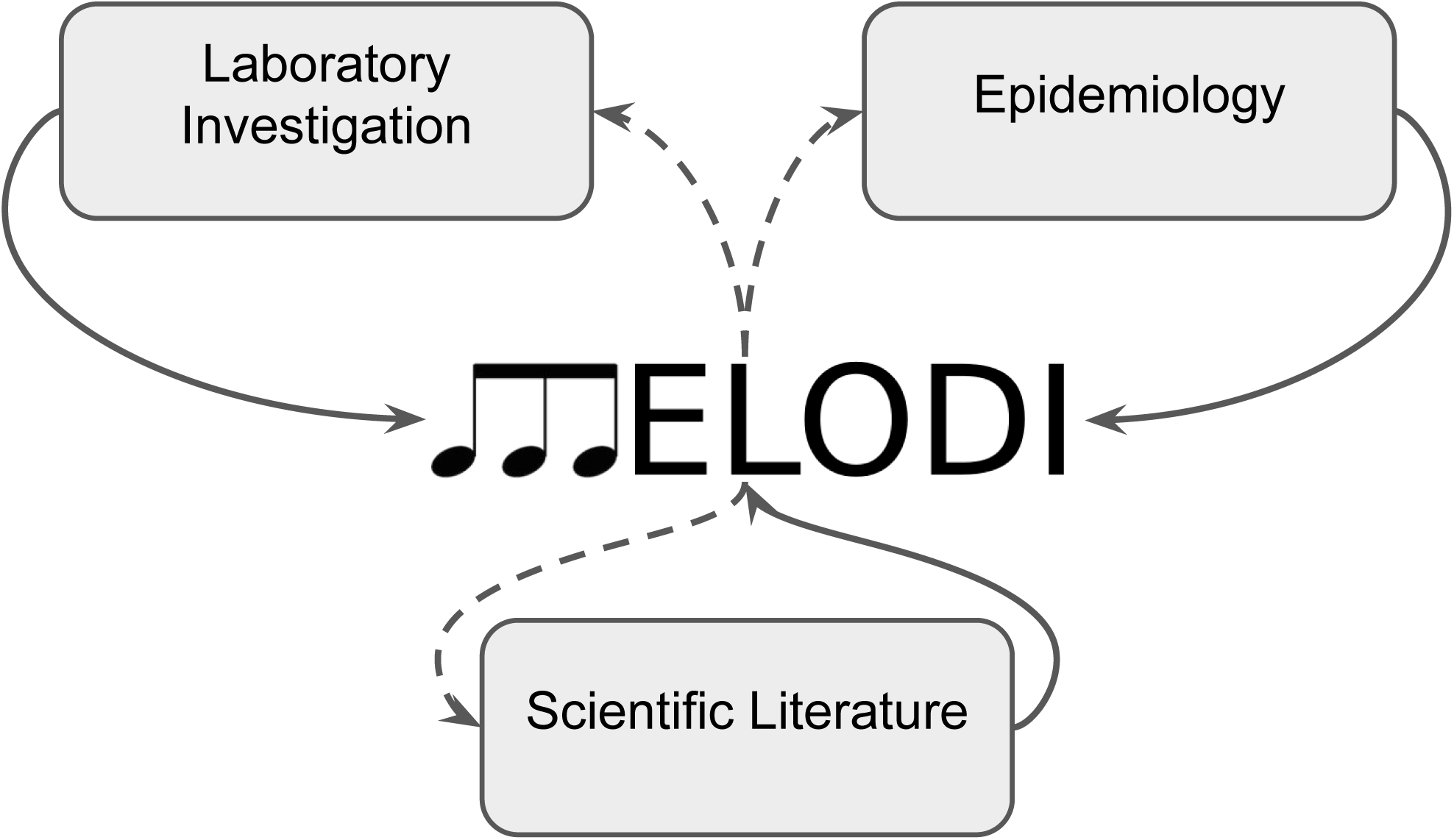
Concept origins and post MELODI investigations. MELODI takes two concepts and derives intermediates. These concepts (solid lines) can originate from a number of places, e.g. an epidemiological observation, or a finding from a laboratory experiment. By creating article sets that represent these two concepts, potential intermediates (dashed lines) can be derived that may provide testable hypotheses. These can then be investigated further using similar approaches to those that created the initial concepts.

#### 1) Laboratory investigation

There are many ways in which laboratory studies could result in two concepts that would benefit from a methodical search of the literature for mechanisms that may connect them. For example, a cell line study that identifies differential levels of a particular protein in a treated cancer cell line compared to untreated could be followed up by a MELODI analysis to identify potential intermediate proteins. These could then be investigated by further laboratory analysis, or through literature review.

#### 2) Epidemiology

Observational epidemiology focuses on broad ‘exposure’ and ‘outcome’ concepts that encompass a lot of underlying mechanisms (using cohort studies and case-control studies in particular). A relationship such as that between alcohol intake and heart disease could be investigated to identify potential intermediate mechanisms that could be investigated in the laboratory, as biomarkers or risk predictors in epidemiology, or via review of the literature. For causal analysis in epidemiology Mendelian randomization (MR) uses genetic variants as proxies(instruments) to investigate whether an exposure (such as alcohol intake) has a causal effect on disease outcome (Smith and Ebrahim, 2001;Timpson *et al*, 2012;Smith and Ebrahim, 2002, 2003). Causal relationships identified with MR could also be investigated in MELODI (exactly as with conventional observational studies), but a real advantage of MR is in following up potential mechanisms identified by MELODI to determine whether they are truly on the causal pathway, or simply biomarkers.

#### 3) Scientific literature

MELODI is based on the scientific literature and can very naturally be used to identify potential mechanisms underlying relationships identified from the literature (particularly those with strong evidence provided by formal systematic review (Egger *et al*, 2008) and metaanalysis). The results of a MELODI analysis are also amenable to literature review for the initial investigation of a potential mechanism. MELODI provides access to all the articles that support a particular mechanism, enabling comprehensive manual review to determine whether the candidate mechanism is plausible. If so, this may be followed up with a formal systematic review or meta-analysis, or using laboratory or epidemiological approaches if additional evidence is needed.

In addition to the approaches described above, an iterative MELODI analysis may be performed, in which an intermediate derived from a MELODI analysis can be used as the starting point for a new investigation using MELODI.

## 3 Results

### 3.1 Performance

We evaluated the time taken for insertion and indexing of all data. This was completed in under two hours and produced a graph database around 50 GB in size and utilising 80 GB RAM. By pre-populating the graph with almost all the necessary data, we greatly reduced the burden of uploading data dynamically. Extracting large amounts of heavily connected data is still, however, a challenge. An article set containing 100,000 articles will take around one minute to create relationships, two minutes for MeSH enrichment and five for SemMedDB enrichment. When comparing two article sets of that size, the overlap step takes two to five minutes, depending on the number of overlapping elements. Therefore, in total, performing a complete analysis on two large article sets can take around ten minutes, depending on server load and queues. However, as the results of each of the enrichment steps are stored to disk they are only performed once. This means that if the same article set is used for subsequent analysis this step is skipped, reducing computational requirements by around 80%.

Our inclusion of a task management system allows multiple users to work with MELODI simultaneously as jobs are either added to an available worker or held in a queue until one becomes available. Currently MELODI is running on a virtual machine with four CPUs (and therefore four workers) but can be scaled easily to support demand by adding more CPUs and distributing the graph on a cluster.

### 3.2 Case Studies

Two case studies describing how MELODI can be used to generate hypotheses and how they can be explored further are described below.

#### 3.2.1 ERG and prostate cancer

ETS-related gene *(ERG)* is an oncogene that has, in the last decade, become closely linked with prostate cancer (Adamo and Ladomery, 2016). Chromosomal rearrangements cause ERG and Transmembrane Protease, Serine 2 *(TMPRSS2)* to fuse together, forming an oncogenic fusion gene which then disrupts the ability of stem cells to differentiate into prostate cells, leading to unregulated and unorganized tissue. This gene fusion is the most common type found in prostate cancer and can be identified by overexpression of ERG in prostate carcinomas (Tomlins *et al.*, 2005).

Much work has been done on elucidating the mechanism by which ERG is associated with prostate cancer. We used MELODI, therefore, to assess the literature on *ERG* and prostate cancer. Article sets for ‘TMPRSS2:ERG or ETS-related gene’ and ‘prostate cancer’ were generated and compared (Mar 1st 2017) using 867 and 138,391 articles respectively. As expected, many genes previously described were identified, notably phosphatase and tensin homolog *(PTEN)* and androgen receptor (AR) in the SemMedDB triple results (Table 3, example 1) and ETS Proto-Oncogene 1 *(ETS1)*, ETS variant 1 *(ETV1)* and ETS variant 4 *(ETV4)* in the SemMedDB single term results (Table 3, example 2). In addition, by ordering this latter set of results by those that share the fewest articles, the Sp1 transcription factor gene (SP1) appears, a transcription factor known to bind to many promoters with a wide ranging set of proposed functions (Figure 5). The SemMedDB data used to identify this intermediate gene was version 26 which included publication data up to April 30th 2016. In August 2016, Sharon et *al* published a paper describing the role of SP1 in the regulation of Insulin Like Growth Factor Receptor 1 *(IGFR1)* by the TMPRSS2-EFG fusion gene (Sharon *et al.*, 2016). In this paper they describe a physical interaction between the *EFG* and SP1 transcription factors, identified by co-immunoprecipitation assays. This work demonstrates the kind of further investigation that could have been performed on the basis of identifying SP1 as a potential intermediate and confirms the value of MELODI in identifying novel intermediates.

**Fig. 5.**
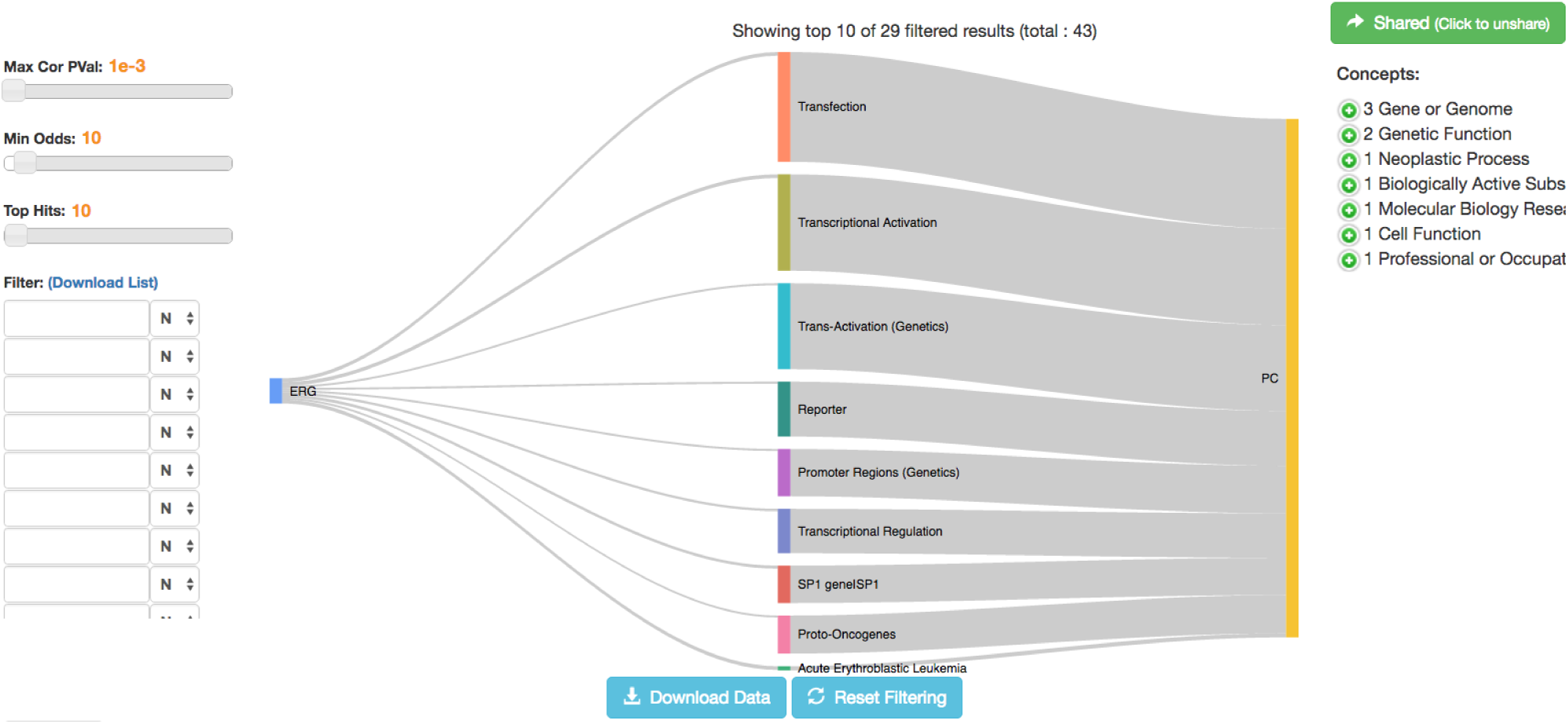
Carnitine and pancreatic cancer. In this example two article sets were compared, one focused around ‘Carnitine’ and the other ‘Pancreatic Cancer’, the results of which are displayed using a force directed graph. Each article set (blue), subject and object (grey) are displayed as nodes with the relationships between them collapsed into single arcs. The thickness of these arcs represents the number of publications generating the relationship.

**Table 3.** MELODI result URLs

| # | URL |
| --- | --- |
| 1 | http://melodi.biocompute.org.uk/results/fedb4912-b04e-4fc8-ac1f-a1c2f04da670/ |
| 2 | http://melodi.biocompute.org.uk/results/2a76380c-324d-4b47-95e4-2dfcaebc5289/ |
| 3 | http://melodi.biocompute.org.uk/results/2f98cf49-a084-4ac3-81ff-3d1932d6bb1d/ |
| 4 | http://melodi.biocompute.org.uk/results/1c5edda6-8fbc-4ea9-b8b6-d66bc82a6a79/ |

In addition, the seven articles supporting the connection between *ERG* and *SP1* are from 2007 and earlier, suggesting the previous studies of Ewing’s sarcoma and acute myeloid leukemia and not prostate cancer may have been overlooked. This indicates that the connection between *ERG, SP1* and prostate cancer could have been identified many years ago. This can be demonstrated by running the same analysis but using publications up to and including 2005, which still finds *SP1* as enriched and overlapping (Table 3, example 3).

#### 3.2.2 Carnitine and pancreatic cancer

The following example illustrates the utility of MELODI in the dissection of causal pathways. Using Mendelian randomization (MR) we have recently found that elevated levels of the amino acid derivative, carnitine, are associated with an increased risk of developing pancreatic cancer (unpublished work at the time of article press). In this situation MELODI can be used as a starting point to investigate how the exposure and the outcome might be connected. Our aim was to generate mechanistic hypotheses that might explain how carnitine increases the risk of pancreatic cancer for further investigation using in vitro studies in the laboratory.

Article sets for ‘carnitine’ and ‘pancreatic cancer’ were created with 14,631 and 82,226 articles respectively and the intermediates derived (Table 3, example 4). Figure 6 shows the enriched relationships identified between these two sets of data and highlights possible intermediates between them. Of interest were the intermediates ‘fatty acid oxidation’ and ‘insulin’. Upon investigation of the literature underpinning these connections (this can be done easily in MELODI which links directly to the articles in pubmed), we found that carnitine can increase fatty acid oxidation (di San Filippo *et al.*, 2003;Lahjouji *et al.*, 2001;Chapoy *et al.*, 1980). Metabolic reprogramming is a known feature of cancer cells (Pavlova and Thompson, 2016) and fatty acid oxidation can be used by cancer cells for energy generation (Rodríguez-Enríquez *et al.*, 2015). We can therefore generate the following hypothesis for further investigation in the laboratory: ‘carnitine increases fatty acid oxidation which provides pancreatic cancer cells with a metabolic advantage’.

**Fig. 6.**
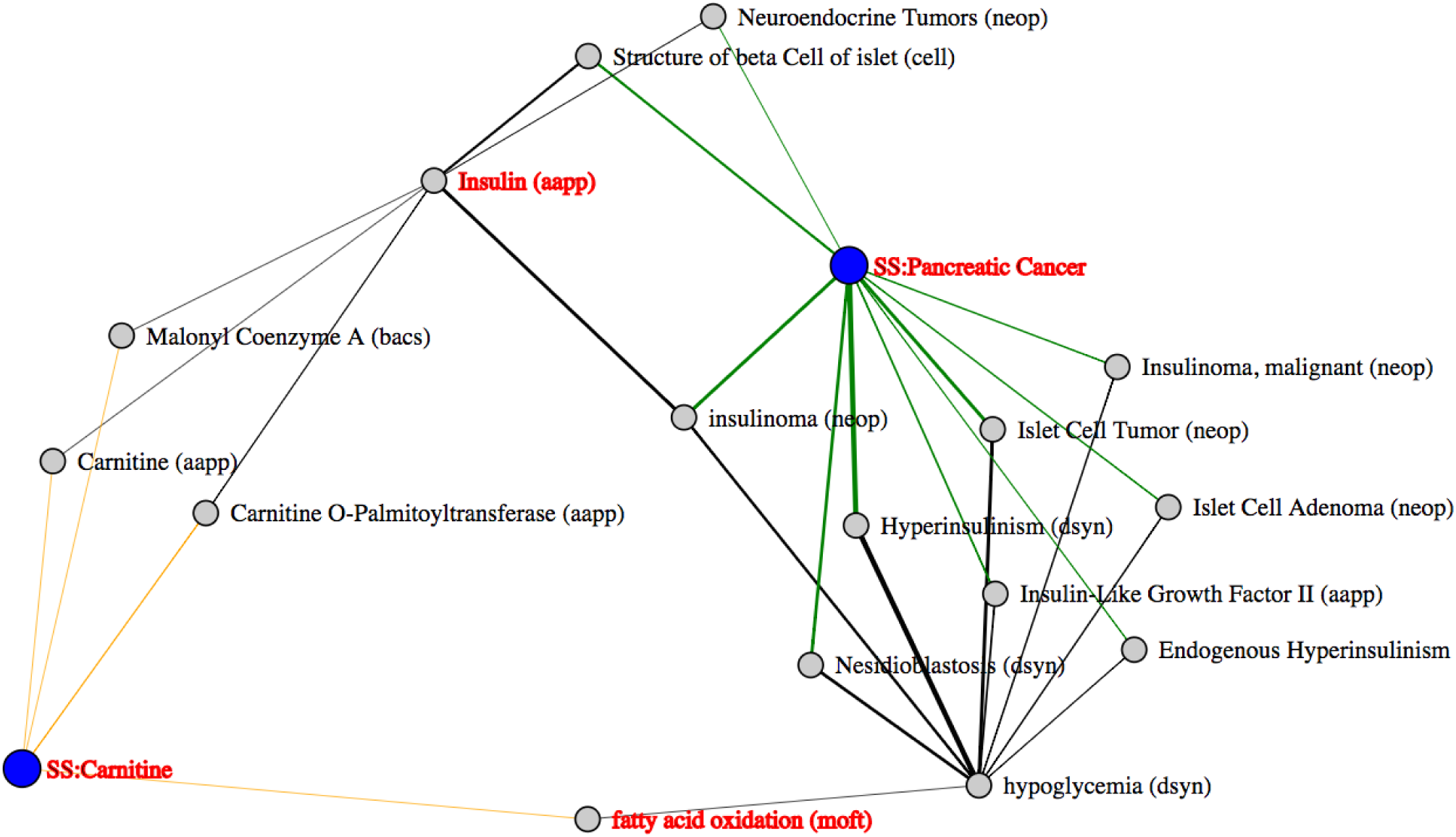
Carnitine and pancreatic cancer. In this example two article sets were compared, one focused around ‘Carnitine’ and the other ‘Pancreatic Cancer’, the results of which are displayed using a force directed graph. Each article set (blue), subject and object (grey) are displayed as nodes with the relationships between them collapsed into single arcs. The thickness of these arcs represents the number of publications generating the relationship.

Insulin is also a highlighted intermediate linking carnitine and pancreatic cancer. Upon investigation of the literature underpinning the connections, we find that insulin treatment of skeletal muscle increases the expression of the carnitine transporter protein OCTN2. Investigation of the literature highlighted by MELODI informs us that increased insulin secretion is associated with some forms of pancreatic cancer (Grygiel *et al.*, 2012). In addition we have also found using MR that elevated fasting levels of insulin are causally associated with pancreatic cancer (unpublished data at time of article press). Therefore with information from MELODI and our own MR investigations we can generate the following hypothesis for further investigation in the laboratory: ‘elevated levels of insulin cause pancreatic cancer in part through increased expression of the carnitine transporter in pancreatic cancer cells’. This demonstrates the power of MELODI as a hypothesis generator to investigate the mechanisms underlying causal relationships.

### 3.3 Limitations

When identifying single overlapping terms the structure of the text is not so important; however, the SemMedDB triple data is dependent on the structure. For example, an article set might contain the triples geneA - ASSOCIATED_WITH - geneB and geneB - ASSOCIATED_WITH - geneA. The underlying directionality provided by SemMedDB will lead to these two triples being treated as separate entities. Cases where predicates imply direction, e.g. INHIBITS and STIMULATES require this restriction but other predicates such as PART_OF and ASSOCIATED_WITH might not. This highlights how the structure of titles and abstracts are key to the extraction of SemMedDB triples and ultimately the identification of overlapping terms. In addition, the assumed directionality from article set A to article set B, and the method for the identification of overlapping SemMedDB triples, will likely miss many intermediates. However, the potential gain of a small number of true positives is not worth the increase in false negatives that would occur if this restriction was removed.

An important limitation of any literature-based tool is that the published literature may be a biased subset, or a biased over-representation, of research that has been undertaken. A large proportion of negative findings are never published, and groups often publish many related papers with similar ideas discussed in the abstract. In addition, the algorithms used to produce the SemMedDB data and the humans used to assign the MeSH terms may introduce unconscious bias. For these reasons, MELODI is always going to be give a biased representation of what is really known about a topic. However, the alternative to a computational approach is manual curation, which is impossible at this scale and potentially prone to much greater bias. As long as the caveats and limitations are understood then the output of this kind of approach can still be valuable and provide reliable hypotheses.

## 4 Discussion

MELODI is a hypothesis-free application that derives mechanisms, both known and novel, from the published literature. It uses a graph database to find enriched relationships between two sets of articles from the entire collection of MEDLINE articles using both the manually curated MeSH terms and computationally derived SemMedDB terms. We have demonstrated its ability to derive both known and unknown intermediates across large complex data sets. Our examples have shown how it can derive novel intermediates for new studies and generate mechanistic targets underlying observed epidemiological associations.

An additional advantage of this kind of approach is the inclusion of data from any MEDLINE article, regardless of impact factor or citation number, which has obvious benefits for the low impact paper. Often, in the course of a scientific investigation, a decision is made to publish when further work could still be carried out, but time or financial constraints are overriding. This has the benefit of releasing information and data early even if the findings and hypotheses have not been validated. The authors of such papers may feel that more could have been done and the resulting work may lack impact, however, tools like MELODI can still utilise the output from these papers and, combined with similar findings, can be used to encourage others to continue this avenue of research rather than the article being lost in the deluge of papers published every day.

MELODI is already capable of deriving reliable intermediate hypotheses. There is, however, potential for future development. The graph database is suited very well to the inclusion of complementary data sets, and the addition of these is planned. They include drug targets, metabolic pathways, and protein-protein interactions. These extra data would permit reliable multi-step connections between article sets using known data connections. The database already contains information on the SemMedDB term types which provides the necessary connections to the extra data and to provide further filtering options, e.g. show only intermediates that are genes, druggable, proteins, etc.

Filtering of the overlapping elements between article sets could also be improved with more computational methods. Machine learning techniques could be applied to train the filtering steps to improve the accuracy and usefulness of intermediates by learning from the user filtering and overlap with well established data sets.

